# Dynamics of mutation accumulation and adaptation during three years of evolution under long-term stationary phase

**DOI:** 10.1101/2020.08.11.246462

**Authors:** Sophia Katz, Sarit Avrani, Meitar Yavneh, Sabrin Hilau, Jonathan Gross, Ruth Hershberg

**Author notes:** Authors contributed equally to this work (Sophia Katz, Sarit Avrani).

## Abstract

Many bacterial species that cannot sporulate, such as the model bacterium *Escherichia coli*, can nevertheless survive for years under resource exhaustion, in a state termed long-term stationary phase (LTSP). Here we describe the dynamics of *E. coli* adaptation during the first three years spent under LTSP. We show that during this time *E. coli* continuously adapts genetically, through the accumulation of mutations. For non-mutator clones, the majority of mutations accumulated appear to be adaptive under LTSP, reflected in an extremely convergent pattern of mutation accumulation. Despite the rapid and convergent manner in which populations adapt under LTSP, they continue to harbor extensive genetic variation. The dynamics of evolution of mutation rates under LTSP are particularly interesting. The emergence of mutators, affects overall mutation accumulation rates as well as the mutational spectra and the ultimate spectrum of adaptive alleles acquired under LTSP. With time, mutators can evolve even higher mutation rates, through the acquisition of additional mutation-rate enhancing mutations. Different mutator and non-mutator clones within a single population and time point can display extreme variation in their mutation rates, resulting in differences in both the dynamics of adaptation and their associated deleterious burdens. Despite these differences, clones that vary greatly in their mutation rates tend to co-exist within their populations for many years, under LTSP.

## Introduction

Evolutionary experiments are designed to study evolution as it occurs. Bacteria such as *Escherichia coli* are particularly useful for such experiments (reviewed in (Barrick and Lenski 2013; Kassen 2014). Bacteria’s great adaptability makes them an excellent model in which to study the dynamics of rapid adaptation. Short generation times enable investigators to rapidly observe bacterial evolution over many generations of growth. The ability to freeze bacterial populations, while maintaining their viability allows one to “go back in time” and compare evolved populations to their ancestors. The relatively small size of bacterial genomes enables cheap and easy sequencing of many clones or population samples from different populations and time points of an experiment. This in turn provides greater resolution in determining the genetic changes associated with the observed evolutionary processes.

Most bacterial evolutionary experiments rely on setups that drive continuous or semi continuous growth and which dilute out non-multiplying cells (Barrick and Lenski 2013; Kassen 2014). This includes both experiments reliant on chemostats and experiments reliant on the serial dilution of evolving populations. Yet, evolution likely often occurs within populations that are not continuously growing and in which the presence of temporarily non-multiplying cells can greatly affect the dynamics of the evolutionary process (Brock 1971; Shoemaker and Lennon 2018). In order to better understand the dynamics of adaptation within such populations it is useful to consider bacterial populations evolving under long-term stationary phase (LTSP). Bacterial species such as *E. coli* that cannot sporulate, can still survive for many decades within spent media (Zambrano et al. 1993; Finkel and Kolter 1999; Finkel 2006; Avrani et al. 2017; Chib et al. 2017). Upon inoculation into fresh media such bacteria will experience a short growth period, followed by a short stationary phase in which cell numbers remain constant. Shortly thereafter bacteria enter a rapid death phase. However, not all cells parish during this death phase. Instead a small fraction of cells enter what is known as long-term stationary phase (LTSP). In LTSP population sizes reduce much more slowly, remaining fairly constant for months and even years. This allows for the maintenance of viability over many years under conditions of resource exhaustion. Mutation accumulation and fluctuations in genotype frequencies observed during the first few months under LTSP suggest that, at least for the few first months, cell replication occurs under LTSP, likely through the recycling of resources (Avrani et al. 2017; Chib et al. 2017).

In July 2015 our lab initiated evolutionary experiments designed to probe the dynamics of adaptation within resource-exhausted populations under LTSP (Avrani et al. 2017). Five independent LTSP populations were established, each by inoculating ~5*10^6^ *E. coli* K12 strain MG1655 cells per ml of Luria broth (LB), for a total volume of 400 ml, within 2 liter aerated flasks. The flasks were placed within an incubator set to 37°C and are shaken at 225 rpm. Other than the addition of water to counteract evaporation, no new external resources were added to these flasks since. These ongoing LTSP populations are sampled periodically and frozen for later sequencing.

The five populations entered LTSP at about day 11 of the experiment. We have previously reported on our analyses of the dynamics of adaptation within these populations over the first four months (127 days) of the experiment (Avrani et al. 2017). Our results showed that, over the first four months under resource exhaustion, LTSP populations genetically adapt through the rapid acquisition of mutations. We found the same genes and sometimes the same specific sites to be mutated across multiple independently evolving populations. Accumulated mutations were very strongly enriched for functional categories, which combined with the high levels of convergence observed suggest that mutation accumulation was governed by strong positive selection (Avrani et al. 2017).

Mutators, deficient in their DNA repair and suffering substantially higher mutation rates than their ancestral genotypes, were shown to frequently arise within lab-evolved microbial populations (e.g. (Sniegowski et al. 1997; Giraud et al. 2001; Voordeckers et al. 2015)). We observed the emergence of such mutatrors in three of our five LTSP populations (Avrani et al. 2017). Mutators were also observed at substantial frequencies within natural bacterial populations and among clinical antibiotic resistant isolates (Gross and Siegel 1981; LeClerc et al. 1996; Mehta et al. 2019). Mutators likely increase in frequencies due to indirect selection in favor of linked adaptations (Wielgoss et al. 2013; Good and Desai 2016; Raynes et al. 2018). At the same time, there is also indirect selection against mutators, due to linked deleterious mutations. Under conditions in which organisms are well adapted to their environment, more mutations will be deleterious than advantageous, leading to selection in favor of reduced mutation rates. Mutators will therefore likely rise to higher frequencies under conditions in which more adaptive mutations are available, or in other words, when a bacterial population is ill adapted to its environment (Wielgoss et al. 2013; Good and Desai 2016; Raynes et al. 2018).

We were surprised to find that even in populations that did not contain any mutators, adaptation under resource exhaustion did not seem to be limited by mutational input. When mutational input is limited, adaptation is expected to occur through hard sweeps (Fogle et al. 2008). Yet, we observed that at every time point of our experiment, up to and including four months several genotypes competed for dominance within each population, in a pattern of clonal interference (Avrani et al. 2017).

In the current study we expanded our analyses of the dynamics of adaptation under resource exhaustion to a much longer time frame of around three years (1095 days). We show that LTSP populations continue to adapt through the highly convergent acquisition of mutations up to three years under resource exhaustion. For non-mutators a majority of mutations appear to be adaptive. Adaptation seems to continue to not be limited by mutational input, as clonal interference is continuously observed. Our new data allows us to further examine the dynamics of mutator evolution. We find that the populations that originally evolved mutators by day 64 of our experiment continue to contain such mutators up to three years under LTSP. In contrast the two populations that did not develop mutators early on, did not develop them later either. We further show that once mutators establish within a population they can evolve into mutators with even higher mutation rates. Cells with very different mutation rates co-exist at high frequencies for years within LTSP populations. Mutators suffering extremely high mutation rates suffer severe deleterious burdens. Despite such burdens they can persist for years under LTSP, alongside cells with much lower mutation rates. Finally, we show that mutator phenotypes can affect not only the overall rates of mutation accumulation, but also the spectrum of mutations and adaptations that occur.

## Results and discussion

### Long-term stationary phase (LTSP) populations maintain constant viability over the first three years under resource exhaustion

The five LTSP populations established in July 2015 were initially sampled daily, then weekly, monthly and later at longer intervals. Each sample was used to estimate viability within its population through the quantification of colony forming units (CFUs) and was then archived by freezing at −80°C. As can be seen in **Figure 1A**, the populations initially grew to ~10^10^ cells per ml, during the first day of the experiment. They then experienced a rapid death phase that slowed down at around day 11, which we consider to be the first day of LTSP. Populations continue to maintain fairly stable viable cell counts at least up to three years (1095 days) under LTSP (**Figure 1A**).

**Figure 1.**
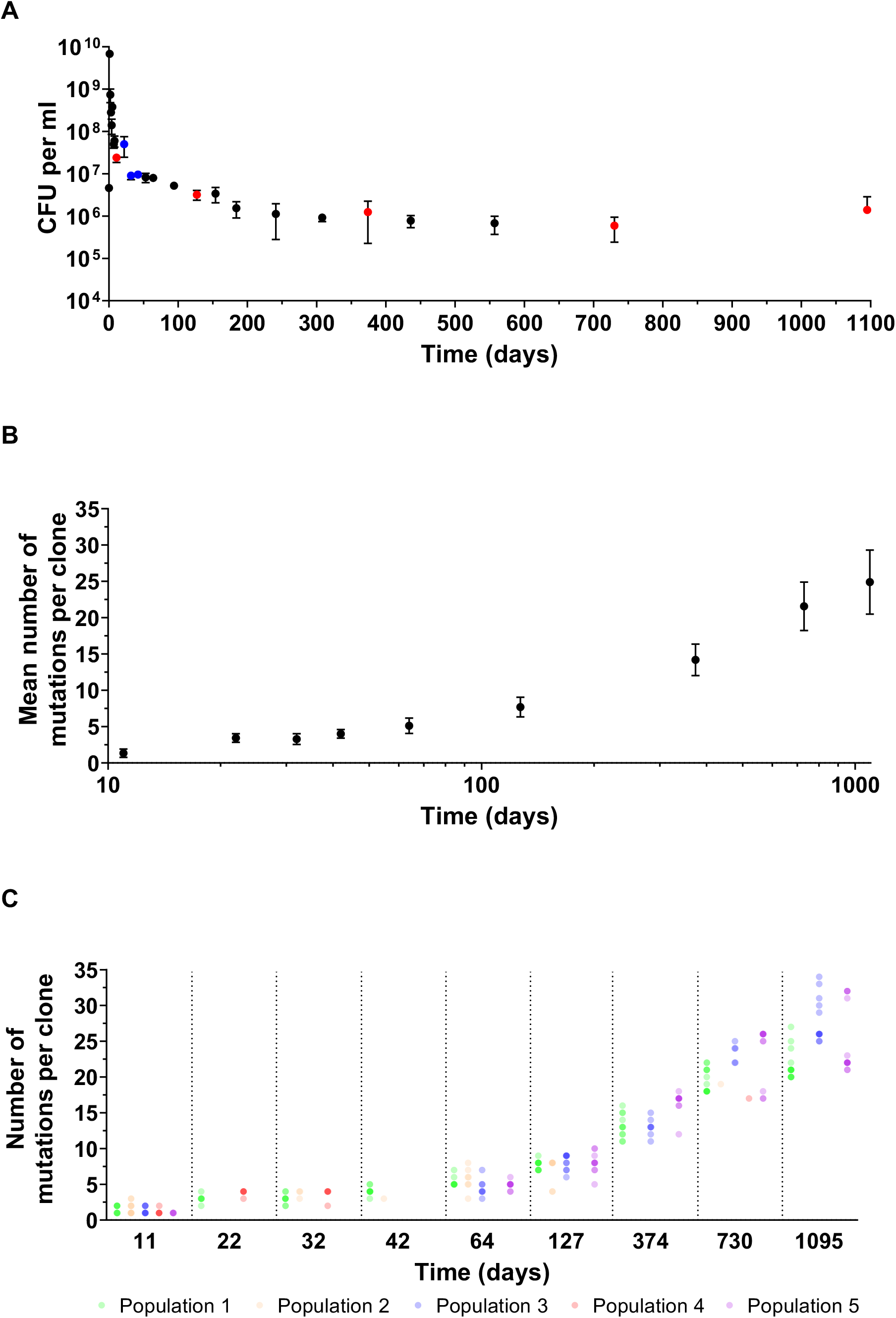
Viable cell counts and mutation accumulation during the first three years under LTSP. (A) *E. coli* populations maintain fairly constant viability levels up to three years under LTSP. Marks represent the mean number of cells per ml of LB, across the five populations as calculated through colony forming unit (CFU) quantification. Error bars represent the standard deviation around this mean. Time points at which ~10 clones per population were sequenced from all five populations are marked red. Time points at which ~10 clones from populations 1, 2 and 4 were sequenced are marked blue. (B) and (C) Continuous mutation accumulation within non-mutator LTSP clones. (B) The mean number of mutations accumulated by LTSP clones increases in a continuous manner across the first three years under LTSP. Error bars represent the standard deviation around the means. (C) Fairly consistent mutation accumulation across populations. Each mark represents an individual clone. Different mark shapes and colors indicate the population from which each clone was extracted. Clones with similar numbers of mutations are represented by overlaid dots, causing a darkening of the dot color.

### *E. coli* LTSP populations continuously adapt in a convergent manner within spent media

We fully sequenced hundreds of clones, sampled from nine time points spanning the first three years spent under LTSP. At each population and time point sampled, ~10 individual clones were sequenced (**Table S1**). The resulting short reads were aligned to the *E. coli* K12 MG1655 genome and mutations were called using the breseq pipeline (Deatherage and Barrick 2014). A full list of mutations found within all clones is given in **Table S2**.

Three of the five LTSP populations include mutator clones, which acquired a mutation within a mismatch repair gene. These clones accumulated a larger number of mutations, compared to non-mutator clones extracted from the same time point and population (**Table S3**, discussed at depth below).

First, we focused on non-mutator clones to examine the rates with which they accumulate mutations with time. As can be seen in **Figure 1B**, non-mutator clones continue to accumulate mutations up to three years under LTSP. Mutation accumulation rates appear to be fairly consistent across populations (**Figure 1C**). We find a significant enrichment in non-synonymous, relative synonymous substitutions (dN/dS >1) for non-mutators across all time points up to and including day 1095 (**Table 1**). Such enrichment in non-synonymous mutations is considered a hallmark of positive selection (Graur and Wen-Hsiung 2000; Ostrow et al. 2014; Tenaillon et al. 2016). Thus, our results show that bacteria are continuously adapting through mutation accumulation, during the first three years under LTSP.

**Table 1.** Enrichment of non-synonymous relative synonymous substitutions in non-mutator clones, up to three years under LTSP

| Time point (days) | # non-synonymous substitutions | # synonymous substitutions | dN/dS<sup>a</sup> | *P*-value<sup>b</sup> |
| --- | --- | --- | --- | --- |
| 11 | 49 | 0 | Can't calculate | <0.0001 |
| 22 | 65 | 0 | Can't calculate | <0.0001 |
| 32 | 38 | 0 | Can't calculate | 0.0006 |
| 42 | 29 | 1 | 8.9 | 0.0091 |
| 64 | 99 | 0 | Can't calculate | <0.0001 |
| 127 | 142 | 8 | 5.5 | <0.0001 |
| 374 | 151 | 23 | 2 | 0.0014 |
| 730 | 258 | 36 | 2.2 | <0.0001 |
| 1095 | 393 | 48 | 2.5 | <0.0001 |

<sup>a</sup>dN/dS (the ratio of the rates of non-synonymous to synonymous mutations) was calculated as
are given in the previous two columns of the table and number of synonymous and non-synonymous sites are calculated based on a combination of all *E. coli* protein-coding genes (see Materials and Methods). dN/dS values could not be calculated for time points in which no synonymous substitutions were observed.
<sup>b</sup> $\chi^2$ P-value with which it is possible to reject the null hypothesis that there is no enrichment in non-synonymous or synonymous mutations, relative random expectations based on the number of non-synonymous and synonymous sites within *E. coli* protein coding genes.

Mutations accumulated up to three years under LTSP accumulate in an extremely convergent manner across the independent LTSP populations. There are 26 genes or intergenic regions, which we found to be mutated within all five populations, 20 mutated in four of the five, and 52 mutated in three of the five (**Table S4**). The loci mutated in 3, 4 or 5 populations constitute only 1.5%, 0.6% and 0.7% of the *E. coli* genome, respectively. Yet, a majority of mutations occurring within non-mutators belonging to each of the five populations fall within these convergently mutated loci (**Figure 2A**). Since convergence constitutes a signal of positive selection (Christin et al. 2010), these results again indicate that the majority of mutations accumulated up to three years under LTSP, within non-mutator clones, are likely adaptive.

**Figure 2.**
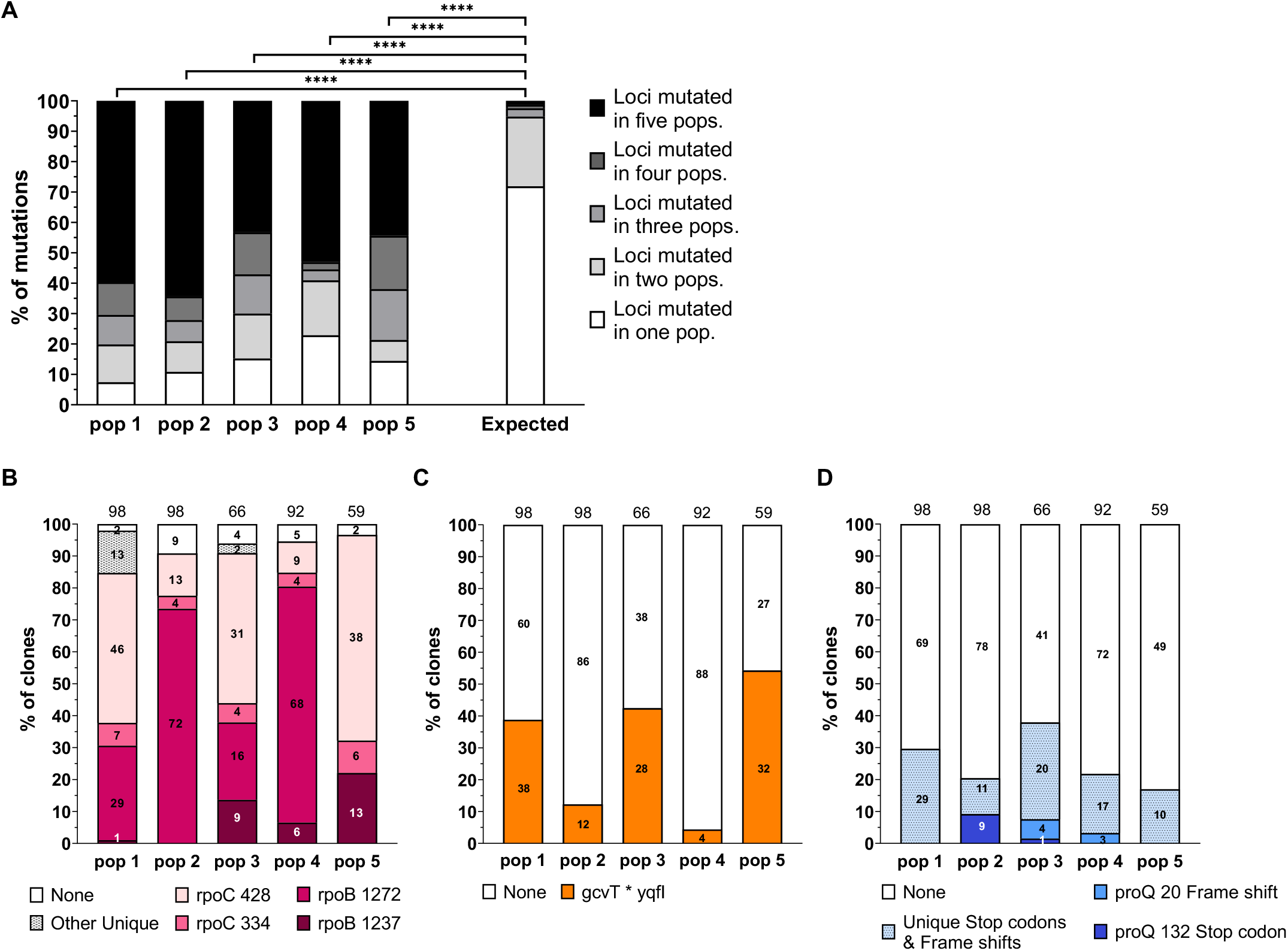
Highly convergent patterns of mutation accumulation under LTSP. (A) The majority of mutations occurring within non-mutator LTSP clones occur within genes that are mutated in a convergent manner. For each population the number of mutations falling within genes mutated in 1, 2,3, 4, or all 5 populations is presented. The relative length of all genes mutated in 1, 2, 3, 4, or all 5 of the populations was used to draw the ‘expected’ bar, which represents random expectations. **** denotes a statistically significant difference (*P* < 0.0001, according to a χ^2^ test) (B) Convergent mutations within the RNA polymerase core enzyme. Depicted is the proportion of clones carrying mutations within each mutated RpoB and RpoC position (for each population). (C) GcvT promoter / 5’ UTR region as an example of loci containing one or at most two specific convergent mutations. All five examples of such loci are presented in Figure S1. (D) ProQ, as an example of genes that are deactivated in a convergent manner across populations. All five examples of such genes are presented in Figure S2. For the (B)-(D), Positions containing a mutation in more than one population are given their own designation. Such positions are marked by the protein name and the position number. If the mutation occurring is a frame shift or stop codon this is indicated. Positions that are mutated in only a single population are grouped within the ‘other-unique’ section of their population’s bar, if they result in a non-synonymous substitution and in the ‘Unique stop codons & frame shift’ section, if they result in a stop codon or frame shift.

The convergently mutated genes include ones encoding some of the most central master regulators of gene expression (e.g. the RNA polymerase core enzyme, RNA polymerase’s housekeeping sigma factor RpoD (σ70), the cAMP-activated global transcriptional regulator CRP, and the translation elongation factor FusA). In addition to being enriched for functions related to the regulation of gene expression, the genes that are mutated in a convergent manner across populations also include many genes related to metabolism and to transport (**Table S4**). We also identified a convergent event that occurred in three of the five populations and involved the deletion of 29 genes, mediated by an insertion sequence (**Table S4**).

### Patterns of convergence are suggestive of whether specific genes undergo changes in their function or loss of function in an adaptive manner

The most striking example of convergence we reported in our previous study involved mutations falling within the RNA polymerase core enzyme (RNAPC). We reported that across all populations ~90% of clones carried a mutation within the RNAPC and that remarkably in the vast majority of cases this mutation fell within one of only three specific sites of the enzyme complex: RpoB position 1272, rpoC position 334 or RpoC position 428 (Avrani et al. 2017). Across the three years of the experiment, the vast majority of clones sequenced continue to carry mutations within the RNAPC that most frequently fall within one of these three sites (**Figure 2B**). A fourth site (RpoB position 1237) is also mutated within four of the five populations (**Figure 2B**). Such convergent changes to specific sites of the RNAPC indicate that specific changes to RNAPC function are adaptive under LTSP.

Interestingly, many clones develop additional mutations within the RNAPC at later time points (**Table S5**). These mutations only occur within clones that already carry one of the four “primary” RNAPC mutations. It is tempting to speculate that these ‘secondary’ RNAPC mutations may be compensatory in nature. In other words, it is possible that the initial RNAPC adaptations carry a beneficial effect on fitness under LTSP, but also incur additional deleterious effects and that the secondary mutations reduce these undesirable effects. A second possibility is that as time progresses under LTSP additional changes to RNAPC function may become advantageous leading to additional mutations within this gene complex. RNAPC mutations were demonstrated to be involved in adaptation to an intriguingly large number of different selective pressures (Conrad et al. 2010; Tenaillon et al. 2012; Avrani et al. 2017; Hershberg 2017). Further studies will be needed to ascertain exactly what these adaptations do to enhance bacterial fitness, under LTSP.

We identified five additional examples in which only one or a few sites of a specific locus were mutated in a highly convergent manner and at very high frequencies across populations (A single example is presented in **Figure 2C**, all five examples are presented in **Figure S1**). Of the five examples, three occurred within the promoter / 5’ UTR regions of genes: A specific mutation within the promoter region of the glycine cleavage system gene *gcvT,* which disrupts the promoter’s known σ70 - 35 hexamer (Santos-Zavaleta et al. 2019) appears at substantial frequencies across all five populations (**Figure 2C**). This is the only mutation seen within the locus, highlighting the likelihood that highly specific changes to the expression of *gcvT* are adaptive under LTSP. The promoter region of the alcohol dehydrogenase gene *adhP* also contains a specific mutation across all five populations (**Figure S1B**). Finally, the promoter / 5’ UTR of the transporter gene *cycA* is mutated at high frequencies across all five populations. In the majority of cases one of two mutations occur within this locus (**Figure S1D**). One of these two mutations appears in four of the populations and is located 43 bases upstream to the gene’s transcriptional start site (Santos-Zavaleta et al. 2019). The second of these mutations appears in three of the five populations and falls within a known binding site of the small RNA GcvB, which regulates the expression of *cycA* at the post transcriptional level (Santos-Zavaleta et al. 2019). Specific high frequency convergent mutations are also observed within the genes *dppA* (**Figure S1C**) and *sstT* (**Figure S1E**). As with RNAPC, the fact that we observe very specific sites to be mutated in a convergent manner across populations indicates that it is specific changes to the expression or function of these five genes that are adaptive under LTSP.

In total we observed 23 specific convergent mutations, each present across three or more of the five independently evolving LTSP populations (**Table S6**).

For five additional genes, we found that across populations deactivating mutations (inserting stops codons or frameshifts) tended to occur (A single example is presented in **Figure 2D**, all five examples are presented in **Figure S2**). These genes were the RNA chaperone gene *proQ* (**Figure 2D**), the transporter genes *glpF* and *dcuA*, the ribosomal protein acetyltransferase gene *rimJ*, and the transcriptional repressor gene *paaX* (**Figure S2**). The convergent occurrence of deactivating mutations within these genes suggests that it may be adaptive to remove the function of these specific genes under LTSP.

### Populations adapting under LTSP maintain high levels of standing genetic variation throughout the first three years of adaptation

Despite the high level of convergence with which adaptations occur between independently evolving LTSP populations, we find that adaptive alleles never fix across any entire individual population. Instead multiple genotypes tend to compete for dominance across all populations and time points (**Figure 3**). As a result, LTSP populations continuously maintain within them very high levels of genetic variation, even as they adapt in a highly convergent manner to prolonged resource exhaustion. Such a pattern of adaptation via soft, rather than hard sweeps indicates that adaptation continues to not be limited by mutational input, even within non-mutators, up to three years under resource exhaustion. Further studies will be required in order to establish whether the maintenance of multiple genotypes over very long periods of time under LTSP is driven by balancing selection, clonal interference, or by other processes.

**Figure 3.**
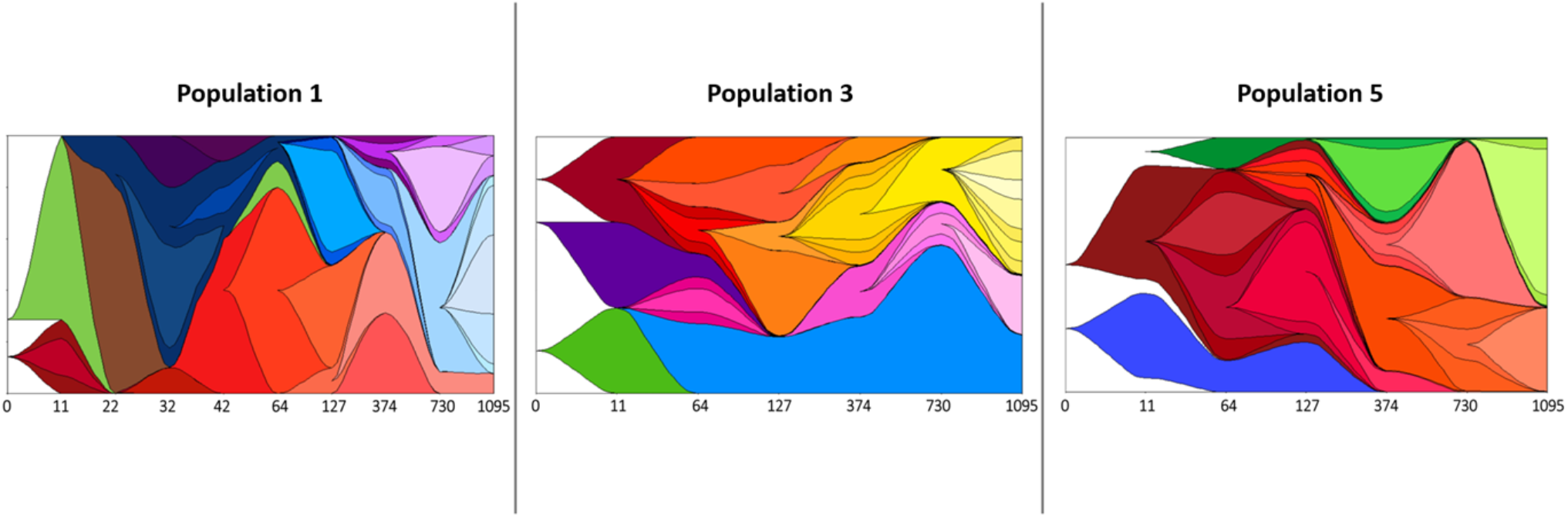
Populations maintain high levels of genetic variation up to three years under LTSP. A clear pattern of clonal interference by which several genotype compete for dominance across time points can be seen. Muller diagrams depicting the relative frequencies of different haplotypes segregating within LTSP populations 1, 3, and 5 are presented. The X-axis indicates the sampling times (not to scale). Only mutations appearing in 30% or more of the considered population’s clones, during at least one time point, were used to generate each plot. In population 3, which included mutator clones once the mutator lineage emerged we do not depict the variation occurring within it, as it is too extensive to draw accurately. This lineage is represented in blue in the population 3 Figure. Due to very high frequencies of mutator clones within populations 2 and 4, we do not present Muller plots for these populations. Muller Plots were produced using the R package MullerPlot (Farahpour et al. 2016).

### Continuous maintenance of variation in mutation rates within mutator containing LTSP populations

As already mentioned, we observed within three of the five LTSP populations the emergence of mutator clones, which acquired a mutation within a mismatch repair gene (**Table S2**). In populations 2 and 3, mutator clones carried mutations within the *mutS* gene. In population 4 a majority of mutators carried a mutation within *mutL*, while a minority carried a mutation within the gene *mutH*. Interestingly, the three populations in which mutator clones were observed by day 64 our experiments, continued to include such clones, at observable frequencies, up to three years into the experiment (**Figure 4A-C**). In contrast, in the two populations in which such clones were not observed by day 64, no such clones were observed up to three years into our experiments. Five populations are not, in our opinion, sufficient to be certain that this pattern by which mutators emerge under LTSP early or not at all is a general trend. However, in the future it might be interesting to examine whether this is indeed a general trend through the establishment of additional LTSP populations.

**Figure 4.**
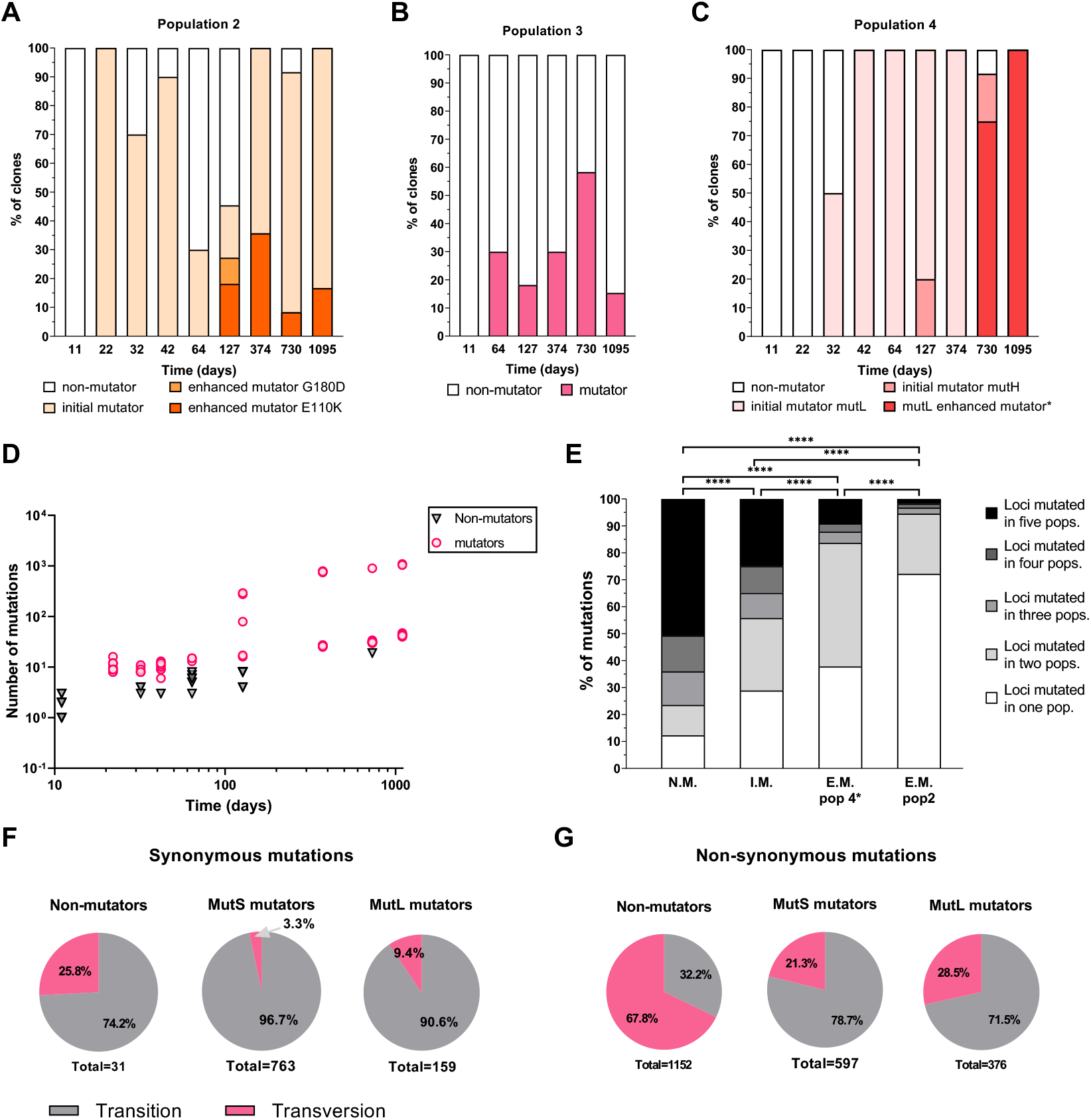
Dynamics of mutator evolution under LTSP. (A)-(C) Clones that vary in their mutation rates co-exist for long periods of time under resource exhaustion in all LTSP populations in which mutators evolved. For each population the relative frequency of each type of clone present within that population is depicted. ^*^In population 4, *mutL* clones sampled from days 730 and 1095 are marked using a different color from *mutL* clones extracted at earlier timepoints, due to their larger than expected mutation accumulation. It is however important to note that we did not observe a significantly higher frequency of development of rifampicin resistance within *mutL* clones extracted during or after day 730 and *mutL* clones extracted prior to day 730. (D) Enhanced mutation rate mutators emerge within population 2. Numbers of mutations accumulated by individual population 2 clones, as a function of time spent under LTSP. Each mark represents an individual clone. Non-mutator clones are represented by black triangles. Mutator clones are represented by magenta dots. Both axis are presented on a logarithmic scale. (E) Higher fractions of mutations fall within convergently mutated genes in clones with lower mutation rates. For each type of clone, the number of mutations falling within genes mutated in 1, 2,3, 4, or all 5 populations is presented. **** denotes a statistically significant difference between bars (*P* < 0.0001, according to a χ^2^ test). (F) Higher transition bias of synonymous mutations, counted only once, within mutators compared to non-mutators. (G) Higher transition bias of non-synonymous substitutions occurring within convergently mutated genes, within mutators compared to non-mutators. For (F) and (G), numbers below each pie chart, indicate the total numbers of mutations used to calculate the percentages presented in that pie chart.

In population 3, mutators and non-mutator clones coexist at high frequencies up to three years under LTSP (**Figure 4B**). In populations 2 and 4, mutator clones rose to very high frequencies, leading to us sequencing only such clones at day 1095 of the experiment. However, we did observe, out of the ~10 clones sequenced, one non-mutator clone in each of these populations at day 730 (**Figures 4A and Figure 4C**). We can therefore conclude that mutator and non-mutator clones tend to co-exist for long periods of time under LTSP. It is important to note that while we did not observe any non-mutator clones in populations 2 and 4 at day 1095, this does not mean they are necessarily not present within the population. After all, by sequencing ~10 clones out of millions of cells present within each population we can only hope to identify the most frequent genotypes.

In population 2, mutator clones, carrying a MutS T300K mutation were first observed at day 22. Up till day 64 clones carrying the T300K MutS mutations had a rather similar number of mutations to each other, which ranged on the order of twice as many mutations as clones that did not carry a mutator mutation (**Table S3**). However, at day 127 mutators seemed to diverge into three clusters (**Figure 4D**). While two of the sequenced mutator clones followed the trajectory of mutation accumulation observed for these mutators so far (17 mutations per clone), three of them accumulated a much higher number of mutations than would be expected from this trend (accumulating 79, 275 and 290 mutations, **Figure 4D**). All mutator clones, irrespective of their accumulated number of mutations, carried the same T300K MutS mismatch repair mutation. We will refer from now on to those mutators carrying a higher number of mutations as ‘enhanced’ mutators. In contrast, mutators carrying a lower number of mutations, consistent with the initial rate of mutations accumulation within mutators will be referred to as ‘initial’ mutators. Within population 2, ‘enhanced’ mutators were also observed at the later time points of our experiments, but always co-existed with ‘initial’ mutators (**Figure 4A,D**).

In order to attempt and identify more of the clones with the ‘enhanced’ mutator phenotype and examine whether these contain a secondary mutation explaining their higher mutation rate, we sequenced five additional clones from population 2 day 127 after plating clones on plates containing the antibiotics rifampicin or nalidixic acid. By sequencing clones that developed antibiotic resistance we hoped to enrich for clones that acquired a larger number of mutations overall. Indeed, all five antibiotic resistant clones sequenced carried the MutS T300K mutator mutation, and each acquired a total of 52, 59, 303, 342 and 609 mutations, placing them in the ‘enhanced’ mutator category. All 16 ‘enhanced’ mutator clones identified throughout the experiment contained a mutation within the gene *dnaQ.* In contrast, none of the 60 ‘initial’ mutator clones carry such a mutation. The *dnaQ* gene encodes the epsilon subunit of DNA polymerase III, which is a 3’-5’ exonuclease responsible for proofreading DNA replication. It was previously shown that mutations within *dnaQ* can greatly increase mutation rates (Echols et al. 1983; Fijalkowska and Schaaper 1996). Intriguingly the three day 127 clones that accumulated between 52-79 mutations carried a G180D DnaQ mutation, while the five that accumulated over 275 mutations by day 127 suffered a different mutation within DnaQ (E110K). At later time points only the E110K mutation was observed, fitting with the higher mutation accumulation rates of all enhanced mutator clones sequenced from day 374 onwards (**Figure 4D**).

Next, we wanted to independently verify that the different types of mutators identified in population 2 indeed differ in their mutation rates, as expected from their patterns of mutation accumulation. To do so, the frequencies with which the four types of clones segregating within population 2 develop resistance to the antibiotic rifampicin (rif), following over-night growth in fresh LB without rifampicin, was compared. As expected, non-mutator clones developed resistance at the lowest average frequency (3.5*10^−9^), compared to ‘initial’ mutators, which developed resistance at a higher average frequency of 5.8*10^−7^. The G180D DnaQ ‘enhanced’ mutators developed resistance at an average frequency higher than that of the ‘initial’ mutators (4.8*10^−6^), but lower than that of the E110K DnaQ ‘enhanced’ mutators (3.7*10^−5^). All reported differences were statistically significant according to a non-paired, one tailed Mann-Whitney test (*P* < 0.01 for all comparisons). Thus, it appears that fitting with rates of mutation accumulation under LTSP, the three mutator types evolving within population 2 indeed vary in their mutation rates, with ‘initial’ mutators, which were the first to emerge having the lowest mutation rates and the E110K DnaQ ‘enhanced’ mutators having the highest.

Different types of mutators also co-exit over time within population 4. However, in this population the differences in mutation rates between the mutator types seem to be less extensive. In population 4, we first observed mutators carrying a mutation within the mismatch repair gene *mutL* at day 32. At day 127 we observed the *mutL* mutators alongside a second type of mutators carrying a mutation within *mutH*. At day 127 both *mutL* mutators and *mutH* mutators carried a rather similar number of mutations (**Figure S3**), which was consistent with the number of mutations observed for the ‘initial’ mutators within populations 2 and 3 at day 127 (**Figure S4**). At day 374 we observed only the *mutL* mutators and these again carried a number of mutations which was consistent with that observed within ‘initial’ mutators within populations 2 and 3 (**Figures S3 and S4**). However, at day 730 we again observed both *mutL* and *mutH* mutators. While the *mutH* mutators seemed to accumulate mutations at a rate consistent with ‘initial’ mutators, the *mutL* mutators acquired more than three times as many mutations, indicating that they may have increased their mutation rate further (**Figure S3**). At day 1095, only the putative ‘enhanced’ mutator *mutL* clones were observed.

To further compare the mutation rates of the different types of mutators found within population 4, we characterized the frequency with which four types of population 4 clones develop resistance to the antibiotic rifampicin (rif) following overnight growth in the absence of that antibiotic. We found that a non-mutator clone extracted from population 4 at day 730 developed resistance to rifampicin at an average frequency of 5.9*10^−10^. The *mutH* clone extracted from the same population and time point developed resistance to rif at significantly higher frequencies (2.4*10^−7^ on average, *P* < 0.001, according to a non-paired one tailed Mann Whitney test). While the *mutH* clone used for this analysis accumulated 31 mutations by day 730, the *mutL* clone we used, which was extracted from the same time point, accumulated 114 mutations. Fitting with this, the *mutL* clone developed rifampicin resistance following overnight growth at significantly higher frequencies, compared the *mutH* mutator clone (3.8*10^−6^, *P* < 0.001). Based on the relative jump in the numbers of mutations accumulated by the *mutL* mutator clones, between days 374 and 730, we expected that the *mutL* clones may have developed a higher mutation rate during that time. However, no significant difference was found between the frequencies with which the day 374 and a day 730 *mutL* clones we examined developed resistance to rifampicin (*P* = 0.215).

### Despite their acquiring a substantial deleterious burden, clones with very high mutation rates can persist for prolonged periods of time, alongside clones with lower mutation rates

We find that ratios of the rates of non-synonymous to synonymous substitutions accumulated by non-mutators, initial mutators and population 4 putative enhanced mutators are all significantly higher than one (**Table 2**). This indicates that within these three clone types mutation accumulation tends to be dominated by positive selection. Further supporting this is the high fraction of mutations accumulated within such clones that fall within genes that are mutated across multiple populations (**Figure 4E**). Positive selection appears to be affecting mutation accumulation more strongly within non-mutators than within initial mutators. This is reflected by significantly higher dN/dS values (*P* < 0.0001, according to a χ^2^ test) and by the fact that within non-mutators significantly higher fractions of mutations tend to fall within genes mutated across larger numbers of populations (**Figure 4E**, *P* < 0.0001, according to a χ^2^ test). In sharp contrast, the vast majority of mutations found within population 2 enhanced mutator clones occur within genes that are not mutated in any of the other populations (**Figure 4E**). Furthermore, mutations accumulated by the population 2 enhanced mutators are significantly enriched for synonymous substitutions, relative random expectations (dN/dS < 1, **Table 2**). This constitutes a signal of purifying selection dominating patterns of mutation accumulation (Graur and Wen-Hsiung 2000). In other words, population 2 enhanced mutators acquire a substantial number of deleterious mutations due to their extremely high mutation rates. Indeed, in order to observe such a dN/dS, which is much lower than 1, many cells of this type would have had to be purged from the population by selection. The extremely high mutation rate of the population 2 enhanced mutators therefore seems to impose on them a high deleterious burden. Yet, despite this deleterious burden they persist alongside non-mutators (at least up till day 730) and alongside initial-mutators (at least up to day 1095), within population 2 (**Figure 4A**).

**Table 2.** Enrichment of non-synonymous or synonymous mutations, relative random expectations, within different clone types

| Type of clones | # non-synonymous substitutions | # synonymous substitutions | dN/dS<sup>a</sup> | *P*-value<sup>b</sup> |
| --- | --- | --- | --- | --- |
| Non-mutators | 1224 | 116 | 3.24 | <0.0001 |
| Initial mutators | 1227 | 263 | 1.43 | <0.0001 |
| Population 4 enhanced mutators | 1269 | 331 | 1.18 | 0.0076 |
| Population 2 enhanced mutators | 2894 | 1583 | 0.56 | <0.0001 |

<sup>a</sup>dN/dS (the ratio of the rates of non-synonymous to synonymous mutations) was calculated as
are given in the previous two columns of the table and number of synonymous and non-synonymous sites are calculated based on a combination of all *E. coli* protein-coding genes (see Materials and Methods).
<sup>b</sup> $\chi^2$ P-value with which it is possible to reject the null hypothesis that there is no enrichment in non-synonymous or synonymous mutations, relative random expectations based on the number of non-synonymous and synonymous sites within *E. coli* protein coding genes.

What advantage could population 2 enhanced mutator clones carry that enables them to persist alongside lower mutation rate clones, despite their deleterious burden? One possibility is that the higher mutation rate of these clones may enable them to accumulate higher numbers of adaptive mutations, counterbalancing the deleterious effects of mutation accumulation. As described above, we identified 98 loci that are mutated in a convergent manner across three or more of the five LTSP populations. The mutations observed within our clones that fall within these loci are likely to be adaptive under LTSP. By day 1095 of our experiments non-mutator, and initial mutators accumulated mutations within 17.1%, and 16.2% of these genes, on average, respectively. At the same time the population 2 enhanced mutator clones accumulated mutations within 48.5% of the convergently mutated genes, on average. Thus it appears that while the higher mutation rates of the population 2 enhanced mutators led to a higher accumulation of deleterious mutations, it also enabled these clones to acquire a higher number of adaptive mutations.

A previous study by Gentile et al. showed that genotypes carrying two mutator alleles (similar to our population 2 enhanced mutators) can outcompete genotypes bearing a single mutator allele (similar to our initial mutators) (Gentile et al. 2011). However, to our knowledge we are the first to demonstrate such an event arising spontaneously within the context of an evolutionary experiment. Indeed, the opposite trend is often expected by which following the emergence of a mutator, anti-mutator alleles will tend to shift rates of mutation back down (Couce and Tenaillon 2019). For example, in Richard Lenski’s long-term evolutionary experiment (LTEE), one of the populations was found to evolve a *mutT* mutator phenotype, increasing mutation rates by ~150 fold. This population was later invaded by a *mutY* anti-mutator allele, which decreased mutation rates by ~40-60% (Wielgoss et al. 2013). It was suggested that this occurred because the supply of adaptive mutations reduced once the LTEE populations became better adapted to the conditions imposed on them, which remain constant with time under the LTEE’s experimental design. Indeed, in the LTEE the greatest fitness gains occurred early on, with a later pattern of diminishing returns (Lenski and Travisano 1994). The fact that in our experiment we see mutators with higher mutation rates co-existing at high frequencies alongside those with lower mutation rates may be the result of adaptive mutation availability remaining high up to three years under LTSP. One possibility for why adaptive mutations may continue to be available under LTSP, is that conditions might be changing, requiring cells to continuously adapt to these changes. A second possibility is that conditions remain fairly constant, but because under LTSP a very strict cap is imposed on how many cells can survive within a population, continuous strong competition occurs between cells, leading to continuous rapid adaptation. More research will be needed in order to distinguish between these two possibilities.

### Mutator evolution can affect both mutational spectra and the spectrum of adaptive substitutions

In addition to affecting mutation rates, mutator mutations also have the potential to affect mutational spectra (i.e. which types of mutations will tend to occur more or less frequently). In order to estimate mutational spectra it is necessary to examine mutations accumulated in the absence of selection (Hershberg and Petrov 2010).

However, mutations accumulated under LTSP are strongly affected by selection. We therefore attempted to examine mutational spectra by focusing on synonymous mutations, which are less likely to be subject to strong selection and by considering each synonymous mutation only once, irrespective of the number of clones it appeared in. For non-mutator clones, ~74% of these synonymous mutations were transitions (i.e. mutations from a purine to a purine or from a pyrimidine to a pyrimidine, **Figure 4F**). The transition bias was even more pronounced within mutators, where over 90% of mutations were transitions (**Figure 4F**). This observed enrichment in transition mutations within mutators was previously demonstrated for a *mutL* mutator in a mutation accumulation study (Lee et al. 2012). Such an increased transition bias may be expected to occur within mutators that are defective in their mismatch repair genes. After all, mismatch repair genes are responsible for repairing errors due to deamination, which tends to lead to C/G to T/A transitions (Zell and Fritz 1987).

Due to the structure of the genetic code, transitions are less likely than transversions to lead to non-synonymous substitutions. The increased transition bias of mutators therefore likely decreases their accumulation of protein-disruptive mutations and by doing so may decrease the deleterious burden they experience. This might contribute to the ability of mutators to persist within our populations alongside non-mutator clones. This fits with a recent study that found that differences in mutational biases between different mutators can lead to variation in the proportion of mutations they suffer that lead to disruption in protein sequences. Computer simulations showed that such differences may potentially greatly affect the strength of selection to decrease the mutators’ mutation rates (Couce and Tenaillon 2019). However, it is important to note that at least for the population 2 enhanced mutators we do find evidence that their extremely high mutation rates do lead to a substantial deleterious burden, and that despite this burden they persist for long periods of time, alongside clones with much lower mutation rates.

Since convergence is a strong signal of positive selection, non-synonymous mutations falling within convergently mutated genes are very likely to be adaptive. It is therefore possible to look at such mutations to study the spectra of adaptive substitutions. As expected from the structure of the genetic code, we find that across clone types, non-synonymous substitutions occurring within convergently mutated genes are more frequently the result of transversion mutations, compared to the synonymous substitutions we examined above. At the same time, non-synonymous mutations within convergently mutated genes are enriched for transitions within mutators, compared to non-mutators (**Figure 4G**). This suggests that differences in mutational biases between mutators and non-mutators translate into differences in the types of adaptive substitutions that ultimately occur.

## Conclusions

We show that *E. coli* populations genetically adapt in a continuous manner up to at least three years under LTSP. Patterns of adaptation are remarkably convergent, with large majorities of non-mutator mutations falling within genes that are mutated across most or all independently evolving populations. This in combination with strong enrichment in non-synonymous vs. synonymous mutations within protein coding genes suggests very strong positive selection affecting mutation accumulation under LTSP. Despite this strong selection LTSP populations continue to maintain very high levels of genetic variation under LTSP, with no single lineage fixing at any given time point, up to and including three years into the experiments. This suggests that despite their existing within spent media for years, adaptation of LTSP populations is not limited by mutational input, even within non-mutators.

Within three of the five examined populations, we saw the emergence of mutator clones fairly early on. Once they emerged, mutators were able to co-exist alongside non-mutators for very long periods of time. In two of the three mutator-containing populations we observed the evolution of mutators with substantially different mutation rates that then co-existed over long time frames. This trend was most pronounced in population 2, where the acquisition of secondary mutation rate-enhancing mutation on the background of an existing mismatch repair mutation, led to the emergence of ‘enhanced’ mutators with extremely high mutation rates. These enhanced mutators then co-existed alongside non-mutators and initial mutators over long periods of time, despite the fact that they suffered a high deleterious burden. The deleterious burden suffered by the enhanced mutators may have been counter-balanced by their ability to accumulate a much higher number of LTSP adaptations. Our results demonstrate that under strong selection it is possible to observe the emergence and long-term co-existence of clones with very different mutation rates and associated deleterious burdens.

Finally, we show that in addition to affecting overall mutation accumulation rates, mutator mutations can also affect mutational biases and types of ultimate adaptive substitutions that emerge, as result of these biases. While overall mutation accumulation rates within mutators are substantially higher, the mutations accumulated tend to be enriched for transitions, which have a lower likelihood of disrupting protein sequences. This may enable mutators to decrease their associated deleterious burden.

## Materials and Methods

### LTSP evolutionary experiments

Five LTSP populations were initiated in July 2015, as described in our previous paper describing the first four months of these experiments (Avrani et al. 2017). Briefly, each population was initiated from a separate clone of *E. coli* K12 strain MG1655. ~5*10^6^ cells per ml were inoculated into 400 ml of Luria broth (LB) within a 2 liter polycarbonate breathing flask. The five flasks were placed in an incubator set at 37°C, where they have been shaking ever since at 225 rpm. No new nutrients or resources were added to the cultures with time, except for sterile water that is added to compensate for evaporation every 10-15 days, according to the weight lost by each flask during that time period.

### Sampling LTSP populations and estimating viability

Initially, every day, then every week, then every month and following that at longer intervals (**Figure 1**) one ml of each culture was sampled. Dilutions were plated using a robotic plater to evaluate viability through live counts. Samples were frozen in 50% glycerol at −80°C.

### Sequencing of LTSP clones

Frozen cultures of the each of the populations and time points (days 374, 730 and 1095) were thawed and dilutions were plated and grown over night. ~10 colonies (**Table S1**) from each culture were used to inoculate 4 ml of medium in a test tube and were grown until they reached an OD of ~1. This procedure was followed in order to minimize the number of generations each clone undergoes prior to sequencing and thus minimize the occurrence of mutations during regrowth. 1 ml of the culture was centrifuged at 10,000 g for 5 minutes and the pellet was used for DNA extraction. The remainder of each culture was then archived by freezing in 50% of glycerol at −80°C. DNA was extracted using the Macherey-Nagel NucleoSpin 96 Tissue Kit. Library preparation followed the protocol outlined in (Baym et al. 2015). Sequencing was carried out at the Technion Genome Center using an Illumina HiSeq 2500 machine. Clones were sequenced using paired end 150 bp reads.

The ancestral clones used to initiate the five populations were sequenced as part of our previous study as were ~10 clones from each of the five LTSP populations at days 11, 64 and 127 and ~10 clones from days 22, 32 and 42 from populations 1, 2 and 4 (**Table S1**) (Avrani et al. 2017).

### Calling of mutations

In order to call mutations, the reads obtained for each LTSP clone were aligned to the *E. coli* K12 MG1655 reference genome (accession NC_000913). LTSP clone mutations were then recorded if they appear within an LTSP clone’s genome, but not within the ancestral genome, which was previously sequenced (Avrani et al. 2017). Alignment and mutation calling were carried out using the Breseq platform, which allows for the identification of point mutations, short insertions and deletions, larger deletions, and the creation of new junctions (Deatherage and Barrick 2014).

### Quantifying rifampicin resistance frequencies for individual LTSP clones following overnight growth in fresh LB

Clones were taken from −80°C storage and streaked to isolate an individual random colony. The colony was inoculated into 4 ml of LB for overnight incubation at 37°C with shaking (225 rpm). Following overnight growth, 100 ul of the resulting culture was plated on LB agar plates and on LB agar plates supplemented with 100 μg/ml rifampicin (Sigma-Aldrich) at the appropriate dilutions. Frequencies of rifampicin resistance were calculated by dividing the CFUs grown on rifampicin containing plates by the CFUs grown on plates with no rifampicin added. For each clone tested at least five independent experiments were carried out.

### Calculating numbers of synonymous and non-synonymous sites within the *E. coli* K12 MG1655 genome

DNA sequences of all protein-coding genes of *E. coli* K12 MG1655 were downloaded from the NCBI database. The contribution of each protein-coding site to the count of non-synonymous and synonymous sites was calculated according to the likelihood that mutations to that site would lead to a non-synonymous or a synonymous change. For example, mutations to the third codon position of a 4-fold degenerate codon would be 100% likely to be synonymous. Such a position would therefore add a count of 1 to the number of synonymous sites. In contrast mutations to the third codon position of a 2-fold-degenerate codon will be synonymous for a third of possible mutations and non-synonymous for two thirds of possible mutations. Such positions would therefore add a count of 1/3 to the number of synonymous sites and 2/3 to the number of non-synonymous sites. In such a manner, we could calculate what proportion of sites, across all *E. coli* K12 MG1655 protein-coding genes are non-synonymous (meaning that mutations to those sites would lead to non-synonymous changes) and what proportion are synonymous.

## Acknowledgments

This work was supported by an ISF grant (No. 756/17, to RH) and by the Rappaport Family Institute for Research in the Medical Sciences (to RH). The described work was carried out in the Rachel & Menachem Mendelovitch Evolutionary Process of Mutation & Natural Selection Research Laboratory.

