## Supplementary material for "Dynamics of mutation accumulation and adaptation during three years of evolution under long-term stationary phase": Figures S1-S4

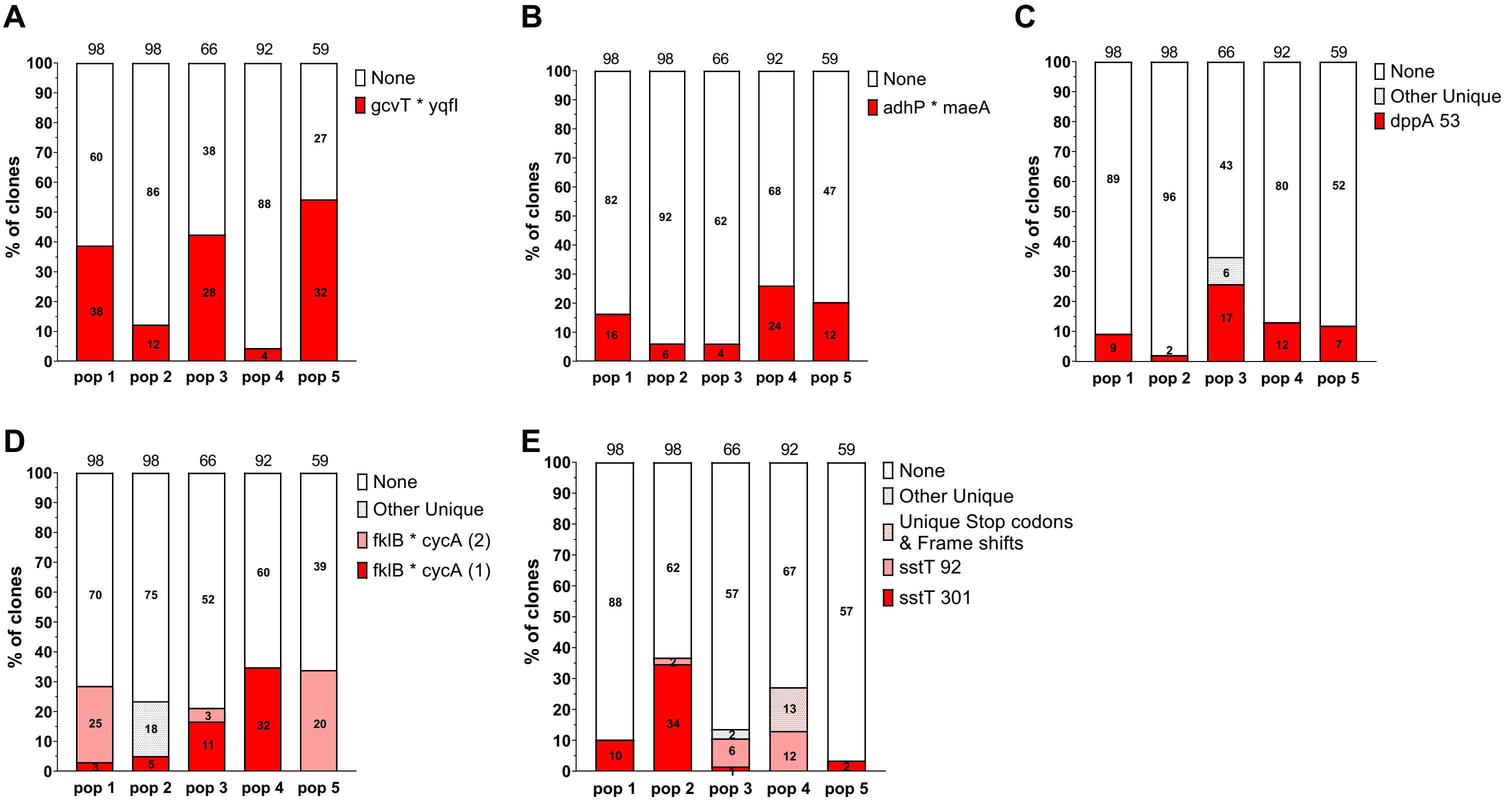


**Figure S1. Loci containing one or at most two specific convergent mutations.** (A) gcvT gene promoter / 5' UTR; (B) adhP gene promoter / 5' UTR; (C) DppA; (D) cycA gene promoter / 5' UTR; (E) SstT. Positions containing a mutation in more than one population are given their own designation. Such positions are marked by the protein name and the position number, if the locos considered is a gene and marked as intergenic if the locus considered is a promoter / 5' UTR region. Positions that are mutated in only a single population are grouped within the 'other-unique' section of their population's bar.

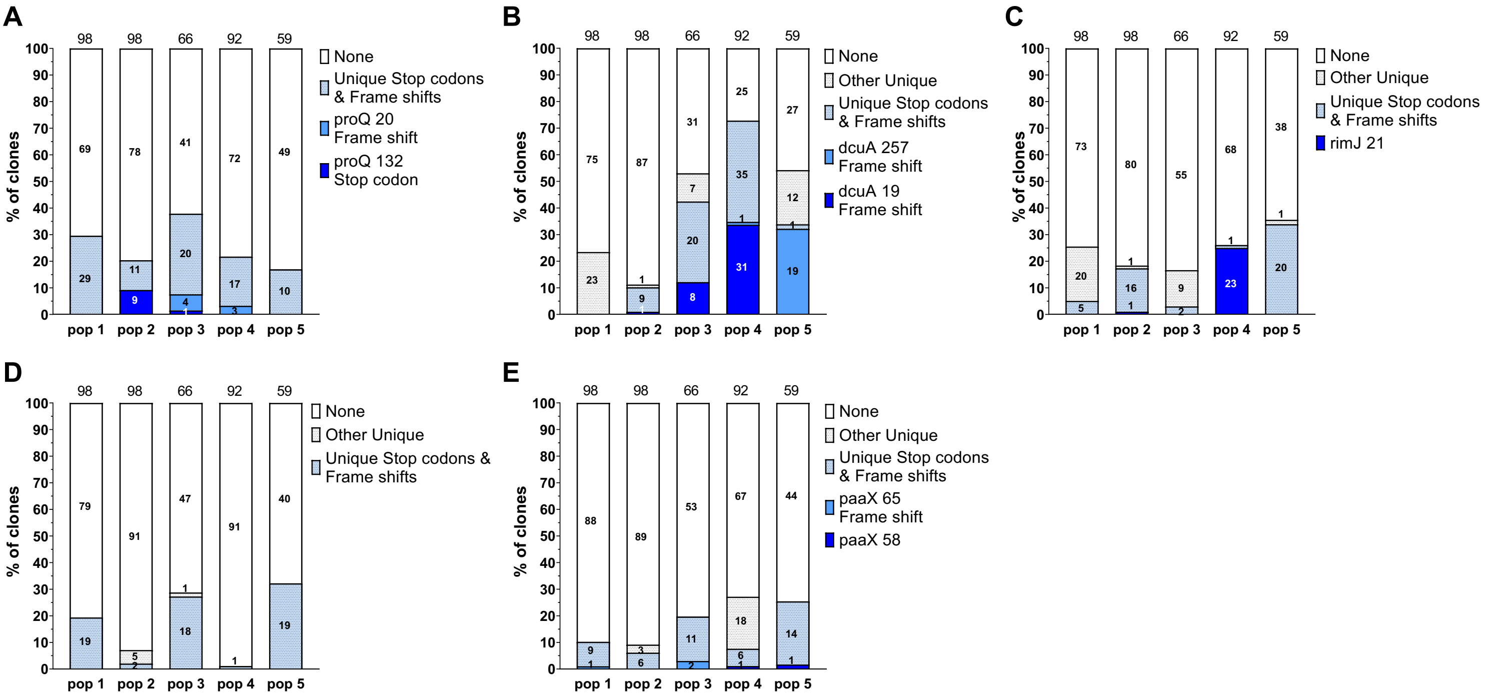


**Figure S2. Genes that are deactivated in a convergent manner across populations.** (A) ProQ; (B) DcuA; (C) RimJ; (D) GlpF; (E) PaaX. Positions containing a mutation in more than one population are given their own designation. Such positions are marked by the protein name and the position number. If the mutation occurring is a frame shift or stop codon this is indicated. Positions that are mutated in only a single population are grouped within the 'other-unique' section of their population's bar, if they result in a non-synonymous substitution and in the Unique stop codons & frame shift section if they result in a stop codon or frame shift.

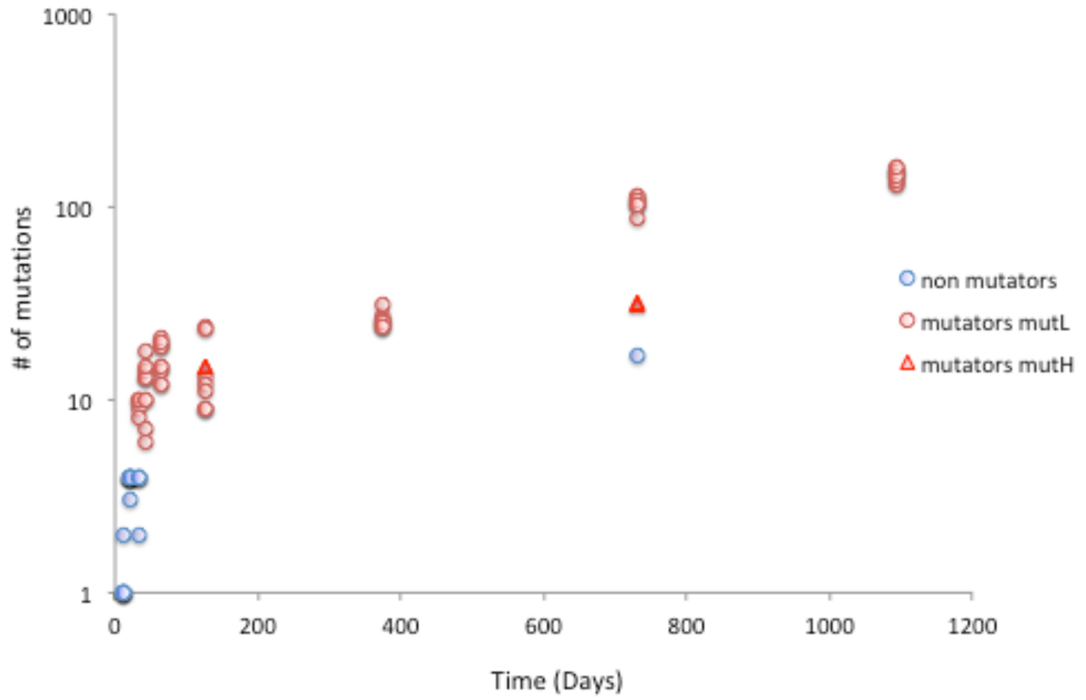


**Figure S3. Numbers of mutations accumulated by individual population 4 clones, as a function of time spent under LTSP**. Each mark represents an individual clone. Non-mutator clones are represented by blue marks. Mutator clones are represented by red marks. The y-axis is presented on a logarithmic scale

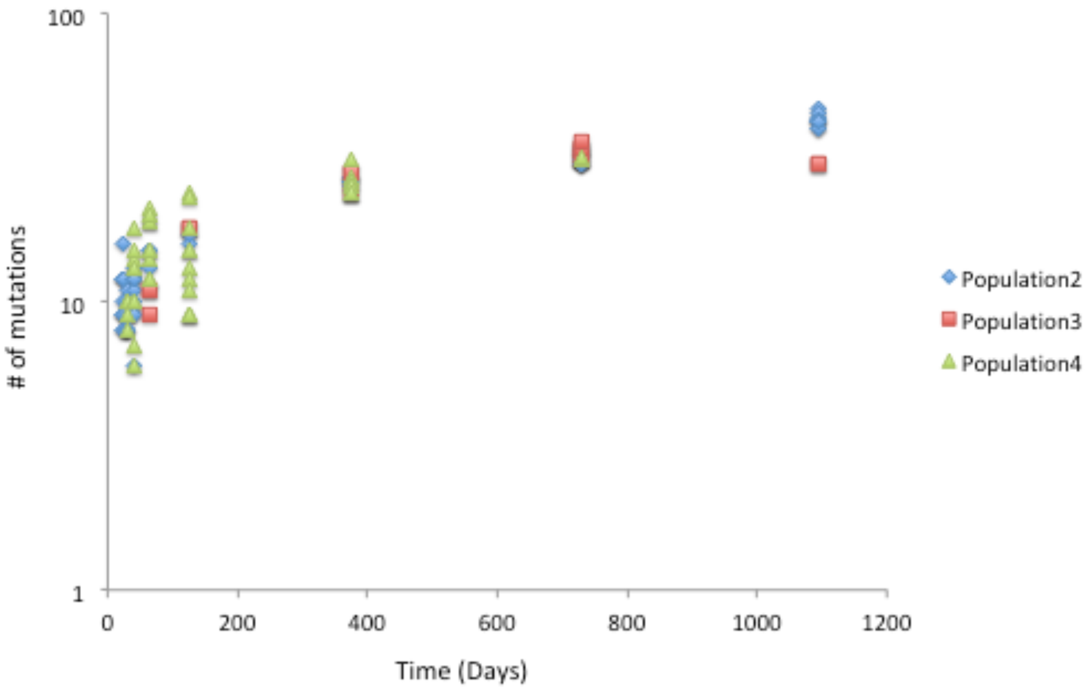


**Figure S4. Numbers of mutations accumulated by individual 'intial-mutator' clones across all three populations are rather consistent.** Marks represent individual clones and are color coded according to the population from which they were extracted. The y-axis is presented on a logarithmic scale.
